# State-Unspecific Modes of Whole-Brain Functional Connectivity Predict Intelligence and Life Outcomes

**DOI:** 10.1101/283846

**Authors:** Yu Takagi, Jun-ichiro Hirayama, Saori C Tanaka

**Affiliations:** ATR Brain Information Communication Research Laboratory Group, Kyoto 619-0288, Japan; Graduate School of Information Science, Nara Institute of Science and Technology, Nara 630-0192, Japan; Japan Society for the Promotion of Science, Tokyo 102-0083, Japan; RIKEN Center for Advanced Intelligence Project, Tokyo 103-0027, Japan

**Keywords:** Functional connectivity fMRI, Machine learning, Human Connectome Project, Intelligence

## Abstract

Recent functional magnetic resonance imaging (fMRI) studies have increasingly revealed potential neural substrates of individual differences in diverse types of brain function and dysfunction. Although most previous studies have been inherently limited to state-specific characterizations of related brain networks and their functions, several recent studies have examined the potential state-unspecific nature of functional brain networks, such as their global similarities across different experimental conditions (i.e., states) including both task and rest. However, no previous studies have carried out direct, systematic characterizations of state-unspecific brain networks, or their functional implications. Here, we quantitatively identified several modes of state-unspecific individual variation in whole-brain functional connectivity patterns, called “Common Neural Modes (CNMs)”, from a large fMRI dataset including eight task/rest states, obtained from the Human Connectome Project. Furthermore, we tested how CNMs account for variability in individual behavioral measures. The results revealed that three CNMs were robustly extracted under various different preprocessing conditions. Each of these CNMs was significantly correlated with different aspects of behavioral measures of both fluid and crystalized intelligence. The three CNMs were also able to predict several life outcomes, such as income and life satisfaction, achieving the highest performance when combined with behavioral intelligence measures as inputs. Our findings highlight the importance of state-unspecific brain networks to characterize fundamental individual variation.

## Introduction

An increasing number of cognitive neuroscience studies have revealed the neural substrates of individual difference using functional magnetic resonance imaging (fMRI) (Dubois and Adolphs, 2016) by investigating coordinated activation (co-activation) patterns of the whole brain. The degree of co-activation between different brain regions of interest (ROIs), often referred to as functional connectivity (FC), is typically measured by the correlation between the blood-oxygen-level-dependent (BOLD) signals averaged within each ROI. A set of brain regions that cooperates under some experimental conditions is typically called a “network”, as represented by the default mode network (DMN) (Raichle, 2015). A wide variety of individual differences in our cognition and behavior have been associated with the characteristics of FC patterns and networks in the brain, including cognitive abilities (Finn et al., 2015; Smith et al., 2015), sustained attention ability (Rosenberg et al., 2016), emotional sensitivity (Modi et al., 2015; Takagi et al., 2018) and psychiatric disorders (Fox and Greicius, 2010; Takagi et al., 2017).

These previous studies have investigated the relationship between individual differences and brain networks while a person is experiencing a specific state. In particular, recent research has intensively focused on the resting state, as it potentially reflects many types of individual differences and can be measured easily (Dubois and Adolphs, 2016). The present study is directly inspired by Smith et al. (2015), who revealed, in a data-driven manner, that a small number of linear factors underlying individuals’ whole-brain resting-state FC patterns (“neural modes”) can explain diverse ranges of individual differences simultaneously (Smith et al., 2015). However, despite their apparent connections with behavior, the brain networks and neural modes examined in these previous studies, as well as their relations to individual differences, are inherently state-specific; thus, it is unclear whether these findings generalize across states, indicating basic traits of individuals. Geerligs et al. (2015) demonstrated that the relationship between individual differences and FC patterns may substantially change across different states, including both rest and task (Geerligs et al., 2015).

A small number of recent studies have suggested the existence of more fundamental, “state-unspecific” brain networks which characterize individuals in a similar manner across different states (Cole et al., 2014; Finn et al., 2015; Tavor et al., 2016). Specifically, Cole et al. (2014) found that average FC patterns of a number of subjects exhibit a high degree of global similarity among different states, including rest. Finn et al. (2015) reported that FC patterns of each individual were also globally similar, across various task and rest states. Furthermore, Tavor et al. (2016) revealed that an individual’s brain activity during a task state can be predicted from their resting-state FC patterns. These findings clearly suggest a potential state-unspecific aspect of brain networks. Unfortunately, however, no previous studies have explicitly identified these state-unspecific brain networks, or quantitatively investigated their relationship to individual differences in behavior.

In the present study, we conducted, for the first time, a quantitative characterization of state-unspecific brain networks and investigated its connection with inter-individual variability in behavior. Our approach combined the large-scale database of the Human Connectome Project with a sophisticated machine learning technique. Specifically, we applied multiset canonical correlation analysis (M-CCA) to the FC matrices obtained from eight states, including the resting state (Kettenring, 1971; Vía et al., 2007). The obtained components uniquely characterize individuals’ FC patterns that are common across different states, which we refer to as “Common Neural Modes (CNMs)”. We demonstrated that several CNMs could be robustly extracted from whole-brain FC patterns. These CNMs were then found to be selectively correlated with behavioral intelligence measures. In addition, we demonstrated that CNMs could predict several types of life outcomes, complementing conventional behavioral measures of intelligence.

## Materials and methods

### Subjects

We used a public fMRI dataset available from the Human Connectome Project (HCP) 500 Subject Release (Van Essen et al., 2012) (http://humanconnectome.org/data). We excluded 1) subjects who did not have all eight fMRI datasets (corresponding to seven task states and one resting state) or who were not given all 44 behavioral measures (subdivided into 12 categories of cognition), and 2) subjects who exhibited substantial movement during fMRI data acquisition (see fMRI preprocessing). After this screening process, 406 subjects were included in the final analysis. All subjects were healthy adults (ages 22–36 years, 238 females).

### MRI parameters

The fMRI data were acquired using a protocol with advanced multiband sequences. Whole-brain echo-planar scans were acquired with a 32-channel head coil on a modified 3T Siemens Skyra with repetition time = 720 ms, echo time = 33.1 ms, flip angle = 52°, bandwidth 2,290 Hz/Px, in-plane field of view = 208 ×180 mm, 72 slices, 2.0 mm isotropic voxels, with a multiband acceleration factor of 8 (Uğurbil et al., 2013). Data were collected over 2 days. On each day, 28 min of rest (eyes open with fixation) fMRI data across two runs were collected (two runs, 56 min in total, per day), followed by 30 min of task-fMRI data collection (60 min in total, per day). Each of the seven task-fMRI was completed over two consecutive fMRI runs. Three task-fMRI (working memory, reward learning, and motor responses) data were collected on the first day. The other four task-fMRI (emotion perception, language processing, relational reasoning, and social cognition) data were collected on the second day. More details about the fMRI collection method were described in previous studies (Barch et al., 2013; Smith et al., 2013).

### Task paradigms

The seven task-fMRI paradigms were selected to activate different neural circuitry that supports broad cognitive functions, and included emotion perception, reward learning, language processing, motor responses, relational reasoning, social cognition, and working memory (Barch et al., 2013; Cole et al., 2016). Briefly, the emotion task involved matching fearful or angry faces to a target face. The reward learning task involved a gambling task involving monetary rewards and losses. The language task involved auditory stimuli consisting of narrative stories and math problems, along with questions to be answered regarding the prior auditory stimuli. The motor task involved movement of the hands, tongue and feet. The relational reasoning task involved higher-order cognitive reasoning regarding relations among features of presented shape stimuli. The social cognition (theory of mind) task used short video clips of moving shapes that interacted in some way or moved randomly, with subjects making decisions about whether the shapes had social interactions. The working memory task involved the conventional visual 2-back and 0-back tasks.

### fMRI preprocessing

Fig. 1 shows a schematic diagram of our analysis. The datasets were originally preprocessed through the HCP minimal preprocessing pipeline (Glasser et al., 2013). This pipeline includes artefact removal, motion correction and registration to standard space. T1 images were segmented into three tissue classes in Montreal Neurological Institute (MNI) space using Statistical Parametric Mapping 8 (SPM8: Wellcome Department of Cognitive Neurology, http://www.fil.ion.ucl.ac.uk/spm/software/) in MATLAB (The MathWorks, Inc., Natick, MA). First, for each subject, the framewise displacement (FD) at each scan was calculated by summing up all six head motion parameters. The “scrubbing” procedure (Power et al., 2012) then identified scans that exhibited excessive head motion based on FD volumes. Specifically, a scan was flagged if the FD exceeded 0.5 mm. The flagged scan, the preceding scan, and the two subsequent scans, were excluded from the correlation analysis below. Subjects were excluded from the subsequent analyses if less than 50% of the scans remained after this procedure for any of the eight fMRI data sets. Then, for each subject, pair-wise, interregional FC was evaluated among 268 ROIs covering the entire brain (Finn et al., 2015) (atlas can be downloaded from https://www.nitrc.org/frs/?group_id=51). The representative time course of each region was extracted by averaging the BOLD time courses of the voxels within that region. Each ROI time course was linearly regressed on the temporal fluctuations of both the white matter and the cerebrospinal fluid as well as the six head motion parameters, whose effects were then subtracted from the original time course. The fluctuation of each tissue class was the average time course of the voxels within the corresponding mask. After within-run linear trend removal, for each subject, we calculated an FC matrix consisting of all the pairwise FCs between the 268 ROIs, based only on the remaining scans after the scrubbing step above. As the FC matrices are symmetric, values on only the strictly lower part were kept, resulting in 35,778 (= 268 × 267 / 2) unique entries (FC values) (Fig. 1a). For all task and resting state fMRI data, FC matrices were calculated using the same procedure. Note that an FC matrix was obtained for every run, and those of multiple runs were averaged in each of the eight task or resting-state conditions.

**Figure 1.**
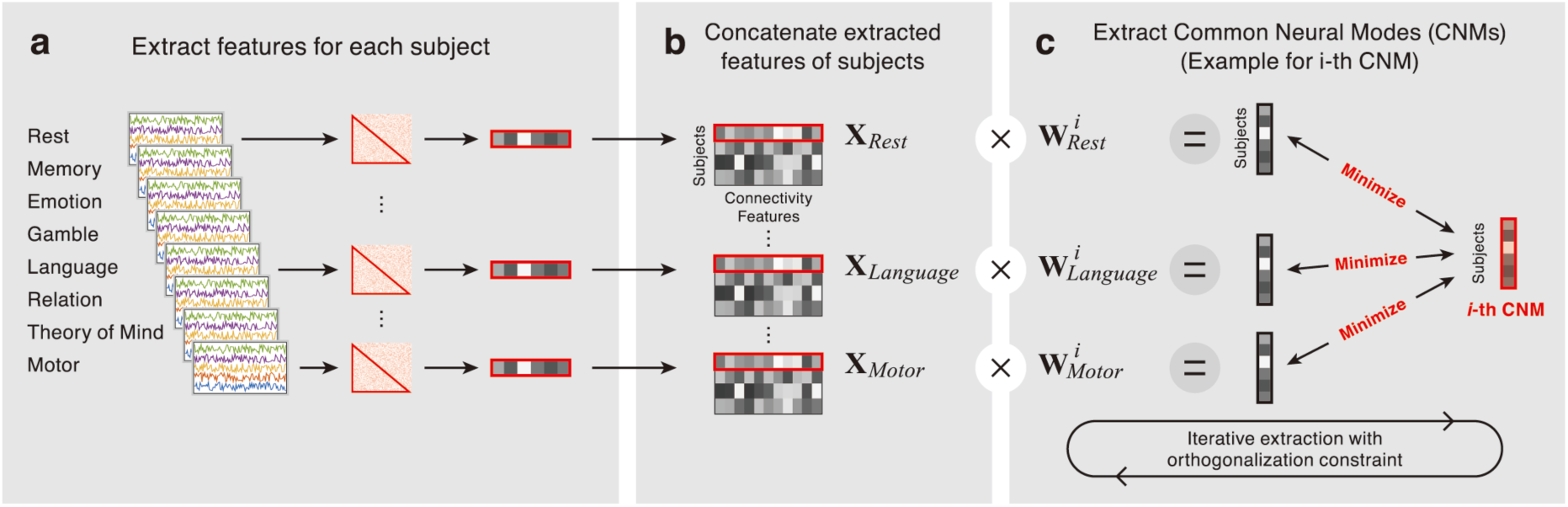
Schematic diagram of the analyses. **(1)** For each subject, feature vectors from the eight states were extracted. (2) Within each state, data for all subjects were concatenated to obtain input data matrices. (3) Common Neural Modes (CNMs) were calculated by minimizing the difference between weighted input matrices and CNMs. CNMs were iteratively calculated with the orthogonalization constraint.

### Identifying CNMs

We identified common neural modes (CNMs) of individuals as FC patterns that robustly characterized individuals irrespective of state. Specifically, we used M-CCA (Kettenring, 1971), which extends canonical correlation analysis (CCA) (Hotelling, 1936) to more than two datasets. Both methods identify canonical variates that summarize each dataset by linear transformations. In contrast, conventional CCA maximizes correlations between a pair of canonical variates, M-CCA maximizes a scalar objective function that summarizes all pairwise correlations among *M* (> 2) canonical variates. M-CCA reduces to CCA when the number of datasets *M* is two. Several variants of M-CCA have been proposed, depending on how it summarizes the pairwise correlations into a single objective function (Kettenring, 1971). We chose the MAXVAR approach because it explicitly introduces common latent factors across different datasets (Vía et al., 2007), which can be naturally interpreted as CNMs.

Suppose that we are given *M* data matrices 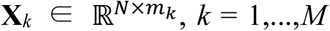 (Fig. 1b), where *N* denotes the sample size and *m*_*k*_ denotes the dimensionality of the *k*-th data space. Each column is assumed to have zero sample mean, without loss of generality. The MAXVAR approach can then be stated as the problem of finding *M* weight vectors **w**_*k*_ (*k* = 1,…,*M*), each for one of the M datasets, so that the errors between the corresponding canonical factors **X**_*k*_**w**_*k*_ and their grand average **z** ∈ ℝ^N×1^ is minimized. The cost function to be minimized is formally given as

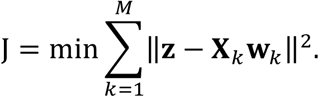

where the minimization is performed with respect to both **w**_*k*_ and **z**. To avoid trivial solutions, **w**_*k*_ and **z**_*k*_ are constrained to have unit Euclidean norms, and to be mutually orthogonal. The solution is given by solving a generalized eigenvalue problem. See Via et al. (2005) for more detailed information about this procedure. Solving this problem gives a set of *M* vectors **w**_*k*_, and CNMs are defined as the average of **X**_*k*_**w**_*k*_ for k = 1,…,*M* (Fig. 1c).

To reduce redundancy among FCs, the dimensionalities of the data matrices were reduced in advance using principal components analysis (PCA). The numbers of principal components were varied between 10, 50 and 100 for calculating CNMs, and the numbers of CNMs were also varied between 10, 50 and 100, respectively. The significance of the pairwise canonical correlations was investigated using a permutation test for individual CNMs. We first shuffled subject labels of all **X**_k_, then conducted M-CCA. We ran these analyses 1,000 times and obtained 1,000 instances of estimated **w**_*k*_. We then took the average of the absolute correlation coefficients between all pairs among **X**_*k*_**w**_*k*_ for each random dataset. Finally, we calculated the statistical significance by comparing the true averaged value of the correlation coefficient with those obtained from shuffled datasets.

### Relationship between CNMs and cognitive measures

To analyze how CNMs were associated with individual differences in behavior, we calculated Pearson’s correlations between the CNMs and cognitive measures obtained using HCP with various behavioral test batteries. The targets of those cognitive measures include, for example, episodic memory, executive function, self-regulation, language and fluid intelligence. The original set of measures were available from the HCP database website. When both age-adjusted and age-unadjusted versions existed for the same index, we excluded the age-unadjusted version.

To reduce the risk of overfitting, we conducted all analyses in a fully cross-validated manner (Barch and Yarkoni, 2013). Specifically, we first split all the subjects into 10 disjointed subsets of subjects. The model for calculating CNMs was then obtained based on all but one set of subjects (training set) and the model was then tested on the one withheld set of subjects (test set). We repeated this procedure 10 times (10-fold cross validation).

### Prediction of life outcomes using CNMs

The preceding analysis suggested that CNMs correlated with representative intelligence measures obtained by the behavioral test batteries. Thus we further investigated whether the CNMs may account for individual differences in the subjects’ life outcomes, which have been considered to be predicted by intelligence measures in the field of educational psychology (Cattell, 1963; Colom et al., 2010; Gottfredson, 1997). As a measure of life outcomes, we chose three measures: income, life satisfaction and year of education. We conducted the analysis using nested 10-fold cross validation. We first split all subjects into 10 sets of subjects, and identified CNMs based on the training set, as with the previous analysis. We then constructed a prediction model using 5-fold cross validation among the training set. We used the L1-reguralized linear regression model for each iteration. The hyper-parameter λ (the regularization coefficient) was tuned by choosing the best value from λ ∈ {0.0001,0.001,0.01,0.1} based on this inner 5-fold cross validation. We finally applied the models for calculating CNMs and life outcomes to the test set. Performance was evaluated by performing Pearson’s correlation between predicted and actual life outcomes across whole subjects.

### Effects of the number of states used to identify CNMs

We investigated the effects of the number of states used to identify CNMs on prediction accuracy. Specifically, we conducted the same prediction analyses as above, but here we used a smaller number of states for constructing the CNMs. We varied the number of states for constructing the CNMs from 2 to 8. We calculated all possible combinations for each case. For example, we calculated 28 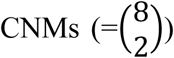, then constructed prediction models for all CNMs, when we estimated the prediction accuracy of two states.

### Interpretation of CNMs

To facilitate the characterization of the biological substrates of the CNMs, we summarized the FC patterns that were correlated with first, second and third CNMs. We focused on these three CNMs because they had been robustly extracted by M-CCA. First, we averaged every FC value over all eight states. We then calculated Pearson’s correlation coefficients between three CNMs and each averaged FC. The 268 ROIs were then grouped into eight representative macroscale networks (e.g., DMN) defined functionally in a previous study (Finn et al., 2015). We then examined the number of FCs between each pair of regions in each network. Finally, we visualized the relative numbers of FCs in each of the two networks as the thickness of the connection lines (see Fig. 5). To aid interpretation, we visualized 200 FCs among all 38,578 FCs that were the most strongly correlated with the CNMs.

**Figure 2.**
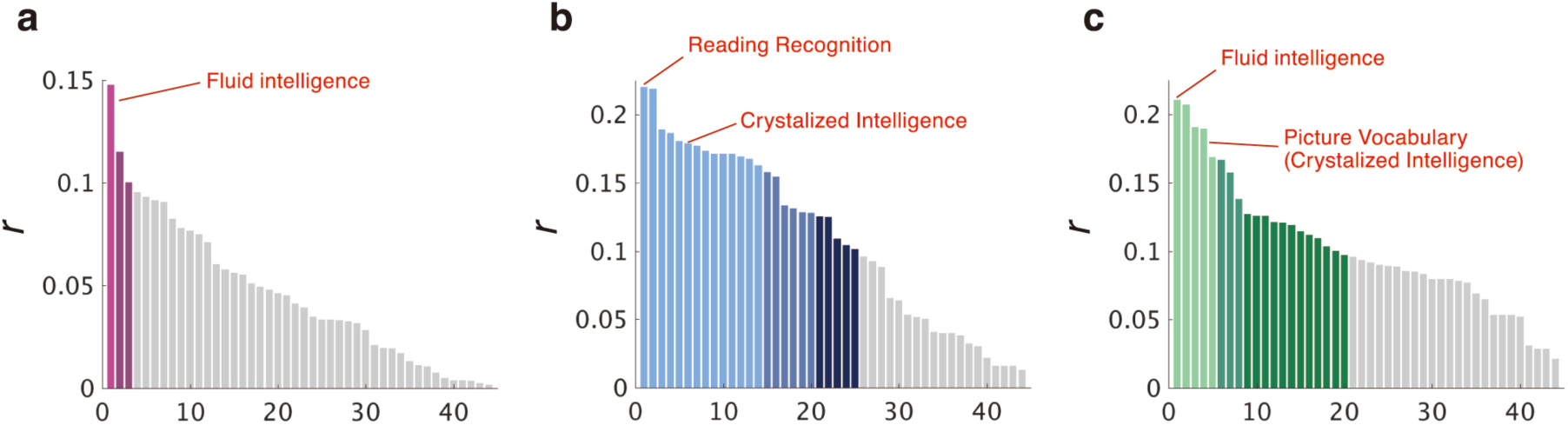
Absolute correlation coefficients (r) between each cognitive measure and the CNMs. Absolute correlation coefficients (r) between 44 cognitive measures and (a) CNM1, (b) CNM2 and (c) CNM3, respectively. The bar with light, medium, dark colored and grey indicated different levels of significance (P < 0.001, P < 0.01, P < 0.05 and P ≥ 0.05, respectively).

**Figure 3.**
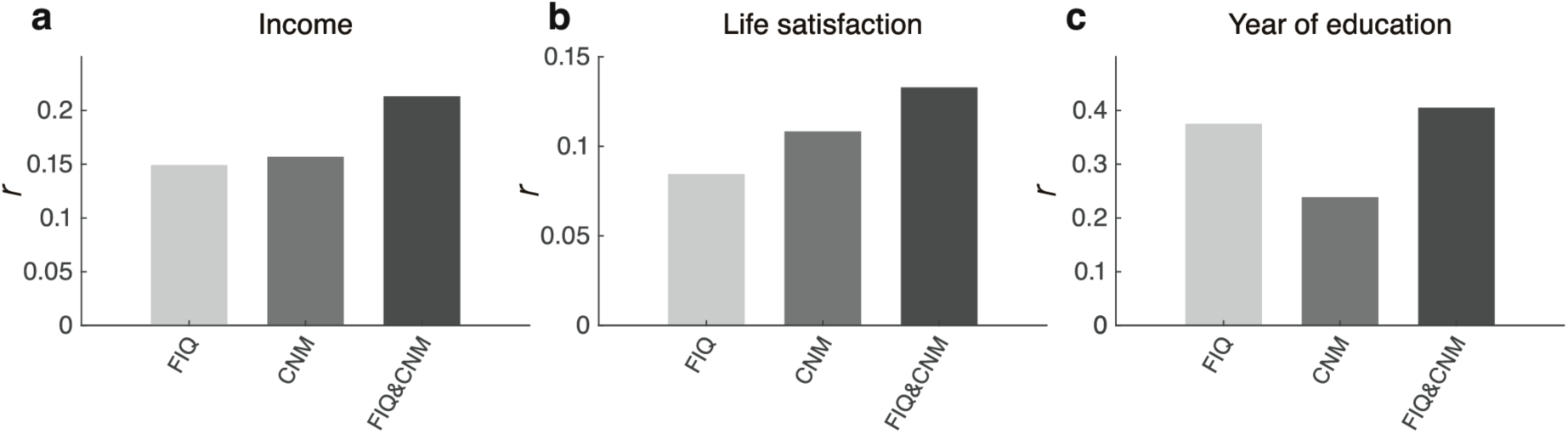
Prediction performance. Cross validated prediction accuracies by the fluid intelligence obtained by the behavioral test batteries (FIQ; left), the CNMs (middle) and their combination (right) for income, life satisfaction and number of years of education, respectively.

**Figure 4.**
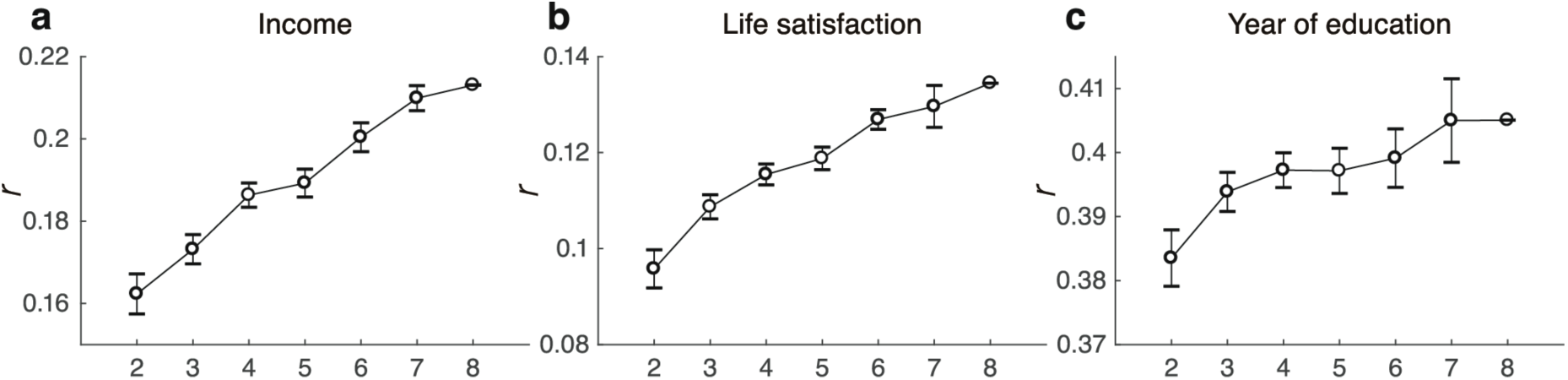
Relationship between the number of states used for the CNMs and prediction performance. Cross validated prediction accuracies of the CNMs obtained from different numbers of states for income (left), life satisfaction (middle) and number of years of education (right), respectively.

**Figure 5.**
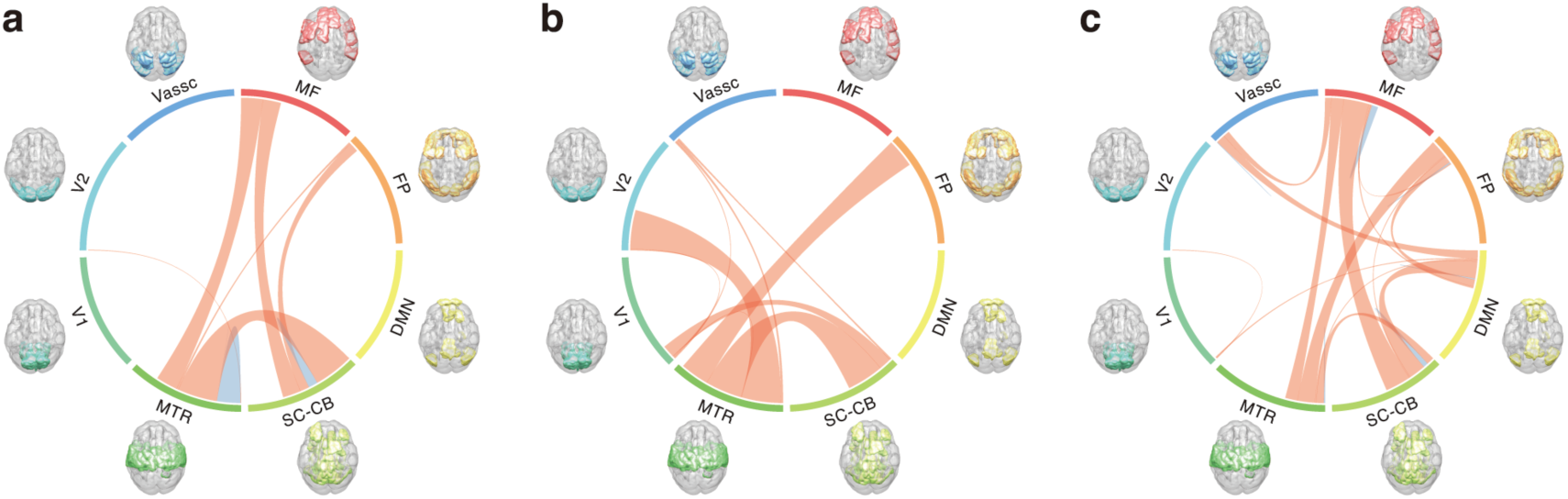
Spatial distribution of the functional connectivity (FC) related to CNMs. The number of FCs between each pair of canonical networks in **(1)** CNM1, **(2)** CNM2 and **(3)** CNM3, respectively. Canonical networks included the medial frontal (MF), frontoparietal (FP), default mode network (DMN), subcortical-cerebellum (SC-CB), motor (MTR), visual I (V1), visual II (V2), and visual association (VAssc). Connection lines are colored blue within the same network and red between two networks.

## Results

### Characterization of CNMs

We first determined the number of CNMs that exhibited significant pairwise canonical correlations among eight states. For any choices of the preprocessing PCA dimensions (i.e., 10, 50, and 100), first, second and third CNMs (namely CNM1, CNM2 and CNM3) exhibited significant overall correlations between states (where all the pairwise correlations were averaged) (*P* < 0.001 for all CNMs; 1,000 times permutation test); the other CNMs did not (*P* > 0.05; 1,000 times permutation test). The M-CCA results were highly consistent under different choices of PCA dimensions (see Supplementary Notes). We therefore focused on the top three CNMs, obtained by M-CCA on 10 PCs of FC vectors.

We then investigated which cognitive measures correlated with each of the three CNMs. Figs. 2a, 2b and 2c show the distributions of the correlation coefficients between cognitive measures and CNM1, CNM2 and CNM3, respectively. Table 1 shows representative behavioral indices with significantly higher correlation coefficients than the chance level. CNM1 was selectively correlated with fluid intelligence, which is a representative component of general intelligence having a broad effect on our daily life and future success (Cattell, 1963; Colom et al., 2010; Gottfredson, 1997). CNM2 correlated with various language related scores (reading recognition and vocabulary comprehension) and self-regulation (delay discounting). It is noteworthy that language related scores are related to crystalized intelligence, a central component of general intelligence along with fluid intelligence (Cattell, 1963; Gottfredson, 1997). Finally, CNM3 was correlated with both fluid intelligence and language-related scores. Note that we confirmed that the correlations described above were not simply the consequence of PCA, which maximizes the variability between individuals in each state (see Supplementary Notes).

**Table 1.**
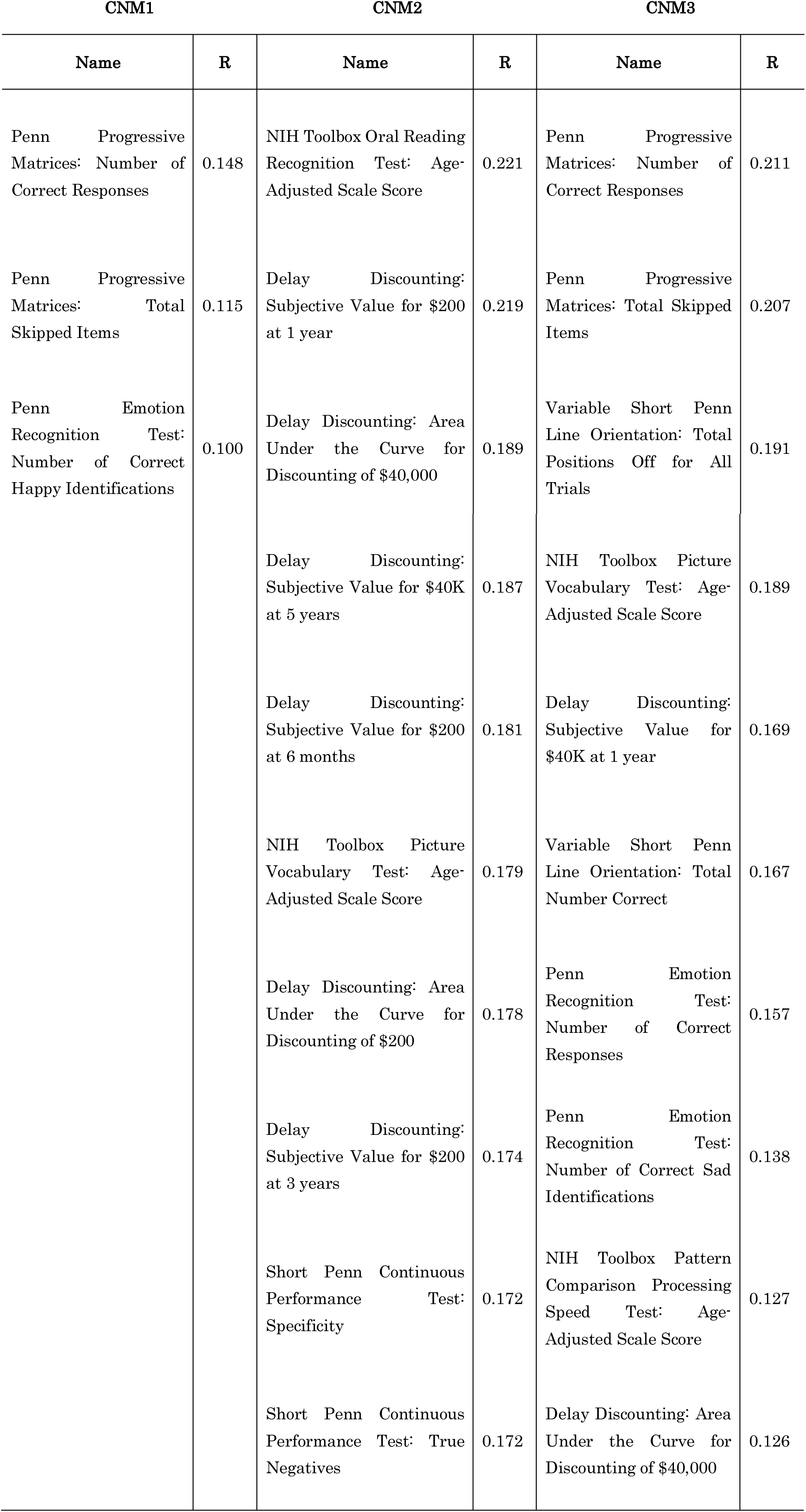
Cognitive measures that were highly significantly correlated with CNMs.

### Prediction of life outcomes using CNMs

We next investigated whether CNMs could predict life outcomes, complementing conventional behavioral tests (i.e., measures of fluid intelligence).

Figs. 3a, 3b and 3c show that predicting with CNMs alone achieved significant predictive value (*P* < 10^-4^ for income and number of years of education; *P* < 2.00 × 10^-4^ for life satisfaction; 10,000 times permutation test). The correlation coefficient (r) was slightly higher than that with fluid intelligence alone for income and life satisfaction, but worse for years of education. Combining both the CNMs and fluid intelligence yielded the highest performance in every case (*P* < 10^-4^ for all income and years of education; *P* < 2.00 × 10^-4^ for life satisfaction; 10,000 times permutation test).

### Effects of the number of states used for the CNMs

We further investigated the effects of the number of states used for identifying the CNMs on the prediction accuracy. Fig. 4a, 4b and 4c show the prediction accuracies using the CNMs with different numbers of states. These figures indicate that the more states we used, the greater accuracy we were able to achieve for predicting life outcomes. We constructed linear regression models, and found that the effects of the number of states were significant for all models (*P* = 8.15 × 10^-13^ for income; *P* = 5.71 × 10^-13^ for life satisfaction; *P* = 0.007 for years of education).

### Interpretation of the CNMs

To facilitate characterization of the biological substrates of the FCs underlying CNMs, we grouped the 268 ROIs into eight macroscale canonical networks. Figure 5 show the circle plots of the FCs that were correlated with CNM1, CNM2 and CNM3. The numbers of FCs in each of the two macroscale regions (the medial frontal [MF], frontoparietal [FP], default mode network [DMN], subcortical-cerebellum [SC-CB], motor [MTR], visual I [V1], visual II [V2], and visual association [VAssc]) networks are presented as the thickness of the connection lines. Connection lines are colored blue within the same network and red between two networks. Although the FCs were widely distributed rather than locally constrained, there were some differences in the distributions among the CNMs. A certain degree of the FCs in the CNM1 belonged to the networks between cortical and subcortical brain regions, including the medial frontal network. On the other hand, FCs in the CNM2 belonged to the networks within cortical brain regions including the frontoparietal network. Finally, FCs in the CNM3 belonged to both the cortico-cortico and cortico-subcortical networks including both the medial frontal and frontoparietal networks.

## Discussion

In the present study, we conducted, for the first time, a quantitative examination of the potential factors underlying state-unspecific inter-individual variability of whole-brain FC patterns, which we termed CNMs, and investigated their associations with behaviors and life outcomes. Although previous studies have suggested a state-unspecific pattern of FC (Cole et al., 2014; Finn et al., 2015; Tavor et al., 2016), to our knowledge no study has directly defined such FC patterns in a quantitative manner. The CNMs were extracted by M-CCA in a fully cross-validated manner from the fMRI datasets of the HCP, covering a broad range of task and resting states. The CNMs predicted representative intelligence measures including fluid and crystalized intelligence with significant correlations, which could not be achieved without M-CCA (i.e., with PCA alone). We further demonstrated that the CNMs were able to predict several life outcomes, complementing conventional behavioral tests of fluid intelligence. We also found that the more states we used to identify CNMs, the higher accuracy we were able to achieve when predicting life outcomes. The FCs constituting those CNMs were widely distributed throughout the brain rather than being locally constrained.

Three CNMs were robustly extracted by M-CCA, which correlated significantly with representative intelligence measures (Fig. 2). Intelligence measures are related to a wide range of cognitive functions and predict broad social outcomes such as educational achievement, job performance, health, and longevity (Cattell, 1963; Colom et al., 2010; Gottfredson, 1997). Therefore, the relationships between the CNMs and these measures are intuitive to understand. It is also noteworthy that each CNM correlated with a different dimension of intelligence. That is, CNM1 and CNM3 correlated with fluid intelligence, while the CNM2 correlated with crystalized intelligence. This suggests that these CNMs may have different biological substrates (Fig. 5). Importantly, the CNMs were derived in a fully data-driven, cross-validated manner. The relationship between CNMs and intelligence measures was thus non-trivial. Although our study was inspired by the “positive-negative” neural modes (Smith et al., 2015) which are also correlated with intelligence measures, our CNM analysis fundamentally differs from that used by Smith et al. (2015) in several important ways. First, although Smith et al. (2015) obtained their results by optimizing the correlation between behavioral measures and FCs explicitly, our CNM did not use any behavioral measure. Second, while Smith et al. (2015) used resting state data only, our CNM method used multiple states.

When predicting life outcomes from CNMs alone, CNMs achieved higher prediction accuracies for income and life satisfaction than prediction with conventional intelligence measures alone. In contrast, conventional intelligence measures achieved better prediction for the number of years of education (Fig. 3). These results may reflect different characteristics between biologically defined measures and measures from a behavioral battery. It should be noted that combining the CNMs with fluid intelligence achieved the highest prediction accuracies for all life outcomes. These results indicate that CNMs contain valuable information for predicting behavior that may not be captured by conventional intelligence measures.

Importantly, using a greater number of states to identify CNMs enabled us to achieve greater prediction accuracy (Fig. 4). This indicates that CNMs were more reliably extracted when considering a greater number of behavioral states. Indeed, the correlation between representative intelligence measures and first principal components derived from each single state were lower than those of the CNMs. Our findings suggest that contrasting many different states, rather than considering any single (typically resting) state, can more reliably identify the neural modes that are able to predict diverse types of individual differences.

Although all three CNMs were related to the subcortical-networks and motor networks, we observed different trends among them in terms of the related canonical networks (Fig. 5). CNM1, CNM2 and CNM3 were related to the medial frontal network, frontoparietal network, and both networks, respectively. This finding is of interest because CNM1 and CNM2 captured different aspects of intelligence (fluid and crystalized intelligence, respectively) while CNM3 was related to both. We also observed that brain regions contributing to all CNMs were widely distributed rather than locally restricted. This is consistent with a previous study reporting that brain regions related to intelligence were broadly distributed (Haier et al., 2009). Several previous studies have also reported a relationship between intelligence measures and FCs (Finn et al., 2015; Lerman-Sinkoff et al., 2017; Schultz and Cole, 2016). However, most of these studies have examined only one state.

Although we focused on state-unspecific neural modes across various states, these modes would be expected to function in a coordinated way with other state-specific neural modes in any particular state. Different neural modes for respective states may have different abilities associated with different neural substrates, which may also cause individual differences in behavior. Thus, it would be useful for future studies to comprehensively compare the relationship between the state-specific and state-unspecific neural modes in terms of their relationship with both cognitive measures and neural substrates.

In summary, we identified neural modes that appeared to be stable across different states, and quantitatively characterized various individual differences. These components, referred to as CNMs, were identified in a fully data-driven manner using a machine learning technique. The CNMs were significantly correlated with representative intelligence measures as well as life outcomes. Although previous studies suggested the potential of brain networks that are shared among broad states, the current study is the first to quantitatively define such networks and demonstrate that they may have a broad effect on behavior and life outcomes. We believe that the present study provides evidence that state-unspecific brain networks may be related to a diverse range of behaviors and life achievements.

## Acknowledgements

This study is the result of “Development of BMI Technologies for Clinical Application” carried out under the Strategic Research Program for Brain Sciences by the MEXT, Japan. JH was supported by KAKENHI 17H06041 from Japan Society for the Promotion of Science (JSPS). We thank Hiroshi Imamizu, Ph.D., Masahiro Yamashita, Ph.D., Ryu Ohata, Ph.D., for helpful discussions and proofreading the manuscript. We thank Mitsuo Kawato, Ph.D., Kazuo Shigemasu, Ph.D., Okito Yamashita, Ph.D., and Ayumu Yamashita for helpful discussions. We thank Nobuyuki Izumihara for his support for visualization.

