## Supplementary Materials for "State-Unspecific Modes of Whole-Brain Functional Connectivity Predict Intelligence and Life Outcomes"

### Supplementary Notes

#### The M-CCA results were robust under different choices of PCA dimensions.

We confirmed that for all pre-selected PCA dimensions, the three CNMs were consistent (for CNM1 and CNM2, $P < 0.001$ for all of the pre-selected PCA dimensions; for CNM3, $P < 0.001$ for the PCA dimensions of 10 and 50, $P < 0.014$ for the PCA dimension of 100; 1,000 times permutation test). The absolute correlation coefficients, averaged over the three PCA dimensions, were 0.92, 0.62 and 0.35 for CNM1, CNM2 and CNM3, respectively ($P < 0.001$ for all CNMs; 1,000 times permutation test).

#### The correlations between CNMs and intelligence measures were not simply the consequence of the representative principal components.

We tested the correlation coefficients between first principal components of each state and fluid intelligence as well as vocabulary comprehension scores. No principal components showed a higher correlation with intelligence measures than correlations between CNMs and intelligence measures. These results indicate that M-CCA uniquely identified state-unspecific characterization of individual FCs, which were specifically correlated with behavioral intelligence measures.
